# Single-shot simultaneous BOLD and velocity-encoding MRI for functional and slow-flow imaging

**DOI:** 10.64898/2026.08.05.743034

**Authors:** Mark Stephan Widmaier, Tzu-Hao Chao, Uzay Emir, Wei-Tang Chang

**Affiliations:** Biomedical Research Imaging Center, University of North Carolina at Chapel Hill, NC, USA; Department of Radiology, University of North Carolina at Chapel Hill, NC, USA; Center for Animal MRI, University of North Carolina at Chapel Hill, Chapel Hill, NC, USA; Department of Neurology, University of North Carolina at Chapel Hill, NC, USA; Biomedical Engineering, University of North Carolina at Chapel Hill, NC, USA

**Author notes:** Mark Stephan Widmaier and Tzu-Hao Chao contributed equally to this work. Corresponding author at: **Department of Radiology**, University of North Carolina at Chapel Hill, NC, USA. E-mail address (W.-T. Chang).

**Keywords:** BOLD fMRI, cerebrospinal fluid, phase-contrast MRI, velocity encoding, BOLD-CSF coupling

## Abstract

Cerebrospinal fluid (CSF) motion is coupled with global blood oxygenation level-dependent (BOLD) fluctuations, but the spatial relationship between regional brain activity and CSF dynamics remains poorly understood. Here, we developed a single-shot BOLD-VENC sequence that combines gradient-echo BOLD imaging with spin-echo velocity encoding following the same RF excitation, enabling simultaneous measurement of brain-wide BOLD activity and spatially resolved slow CSF velocity at 3T. The velocity measurement was validated in a slow-flow phantom and in five healthy participants using paced-breathing, breath-holding, and visual-stimulation experiments. Phantom measurements showed strong agreement with prescribed velocities over 0.1-1.0 mm/s (R^2^ = 0.93-0.98). In vivo measurements demonstrated respiratory- and cardiac-dependent changes in CSF velocity magnitude and direction across the ventricles and cortical subarachnoid spaces (SAS). The established coupling between the negative derivative of the global BOLD signal and fourth-ventricle CSF inflow was reproduced, with a peak lag of 0.9 s. Global BOLD fluctuations were also coupled with spatially distributed CSF velocity changes across ventricular and cortical CSF spaces, with a similar peak lag of 1.2 s. During visual checkerboard stimulation, BOLD-CSF velocity coupling was localized primarily to the SAS surrounding the activated visual cortex, demonstrating a regional relationship between local BOLD activity and nearby CSF motion. These findings establish the feasibility of simultaneous BOLD and slow CSF velocity imaging and extend BOLD-CSF coupling from a global measure toward spatially resolved assessment of hemodynamic-CSF interactions.

## 1. INTRODUCTION

Cerebrospinal fluid (CSF) contributes to central nervous system homeostasis by providing mechanical protection, transporting signaling molecules, and supporting the removal of metabolic waste (Johanson et al., 2008). Within the glymphatic framework, CSF enters the brain along periarterial spaces, exchanges with interstitial fluid, and facilitates the transport of solutes toward perivenous and meningeal drainage pathways (Iliff et al., 2012; Louveau et al., 2015). Although the relative contributions of convection, dispersion, and diffusion to brain-wide solute transport remain under investigation, altered CSF and glymphatic dynamics have been associated with aging (Da Mesquita et al., 2018; Kress et al., 2014) and several neurological conditions, including Alzheimer’s disease (Da Mesquita et al., 2018; Han et al., 2021), Parkinson’s disease (Ding et al., 2021), traumatic brain injury (Iliff et al., 2014), and cerebral small vessel disease (Zhang et al., 2021). Increased extra-axial CSF volume has also been reported in infants who later developed autism spectrum disorder (Shen et al., 2017). Moreover, reduced coupling between CSF inflow and global brain activity in humans has been associated with greater cortical amyloid burden, subsequent cognitive decline, and increasing severity of Alzheimer’s disease-related pathology (Han et al., 2021). Understanding the physiological mechanisms that regulate CSF movement is therefore important for clarifying its potential role in brain health and neurological disease.

Recent functional MRI (fMRI) studies have demonstrated coupling between low-frequency global blood oxygenation level-dependent (BOLD) fluctuations and CSF inflow (Fultz et al., 2019; Han et al., 2021). MRI assessment of CSF dynamics currently relies on several acquisition strategies, each providing different measurement sensitivities. Inflow-sensitive fMRI measures signal changes produced by unsaturated CSF entering the imaging volume, typically near the fourth ventricle or the craniospinal junction, and has been widely used to characterize the temporal coupling between BOLD fluctuations and CSF inflow. Phase-contrast MRI provides a direct measure of velocity by applying bipolar velocity-encoding (VENC) gradients, which generate a phase shift proportional to motion along the encoding direction (Lotz et al., 2002; Pelc et al., 1991).

Gradient-echo phase-contrast MRI is commonly used to quantify relatively rapid, pulsatile CSF flow in the cerebral aqueduct, foramen magnum, and spinal canal (Korbecki et al., 2019). For slower CSF motion within the ventricles and subarachnoid space (SAS), spin-echo phase-contrast acquisitions with low VENC values can provide sensitivity to submillimeter-per-second velocities and reduce contributions from rapidly flowing intravascular spins (Dong et al., 2025). In addition, labeling- and tracer-based MRI approaches have provided complementary measures of CSF transport, exchange, and clearance over longer temporal scales (Ringstad et al., 2018; Yamada et al., 2008).

Although these imaging methods provide complementary information about CSF dynamics, the relationship between regional brain activity and regional CSF motion remains poorly understood. Inflow-sensitive fMRI has established a temporal relationship between global BOLD or cerebral blood flow (CBF) fluctuations and CSF inflow at the fourth ventricle or craniospinal junction (Fultz et al., 2019; Im et al., 2025). However, because the inflow signal is measured only at the imaging boundary, it does not provide spatially resolved information about CSF motion throughout the brain. In contrast, phase-contrast MRI provides measurements of CSF flow velocity and direction with spatial localization. However, BOLD fMRI and phase-contrast MRI are generally acquired in separate scans, each optimized for its respective contrast. Consequently, regional brain activity and nearby CSF motion cannot be measured under exactly the same physiological and motion conditions. Simultaneous acquisition of BOLD and CSF velocity would enable direct investigation of whether localized neural activity is associated with changes in CSF velocity or temporal delay in the surrounding SAS.

To address this need, we developed a single-shot BOLD-VENC sequence that combines a gradient-echo BOLD readout with a spin-echo phase-contrast readout following the same RF excitation. A gradient-echo echo-planar imaging readout first measures the BOLD signal, after which the gradient moments are rewound before a velocity-encoding preparation and a spin-echo echo-planar readout. The relatively long spin-echo echo time increases sensitivity to slowly moving CSF while suppressing signal contributions from intravascular blood because the T2 of CSF is substantially longer than that of blood (Spijkerman et al., 2018; Zhao et al., 2007). Our BOLD-VENC sequence provides brain-wide coverage and can encode velocity along each of the three orthogonal spatial directions, producing intrinsically registered BOLD and CSF velocity measurements within the same acquisition. In addition, when the inferior imaging slice is positioned at the fourth ventricle or another region dominated by CSF inflow, inflow-sensitive CSF signals can also be measured provided that the repetition time (TR) is shorter than the T1 relaxation time of CSF. This capability enables direct comparison between inflow-sensitive and velocity-encoded measurements of CSF dynamics within a single acquisition. We evaluated the proposed method using controlled flow-phantom experiments to assess the accuracy and sensitivity of the sequence at the velocities in submillimeter-per-second range. For the in vivo experiments, we employed paced breathing and breath-hold tasks to determine whether the sequence could detect physiologically expected changes in intracranial CSF flow magnitude and direction. Given that the respiratory tasks primarily modulated BOLD activity across the brain, we employed visual stimulation to produce localized BOLD activation and investigate its relationship with CSF velocity in the surrounding subarachnoid space. Together, these experiments demonstrate the feasibility of simultaneously measurement of brain-wide BOLD activity and slow CSF velocity, providing a new approach for investigating their spatially resolved relationships.

## 2. MATERIALS AND METHODS

### 2.1 BOLD-VENC sequence

The proposed BOLD-VENC sequence consists of three modules: a gradient-echo (GE) echo-planar-imaging (EPI) readout, a VENC preparation module, and a spin-echo (SE) EPI readout as shown in Figure 1. The imaging parameters were a spatial resolution of 2 mm isotropic and a field of view (FOV) of 192 × 192 × 96 mm^3^. The gradient-echo module acquired the BOLD signal with an echo time (TE) of 33 ms. All gradients were rewound to zero after GE EPI readout. The VENC was set to 1.6 mm/s. The spin-echo EPI readout used a TE of 189 ms. The long TE makes VENC module sensitive to slow bulk flow while maintains low b value (49 s/mm^2^) to minimize signal attenuation. Additionally, the intravascular signal is expected to be largely suppressed, resulting in minimal vascular contribution to the measured signal.

**Figure 1:**
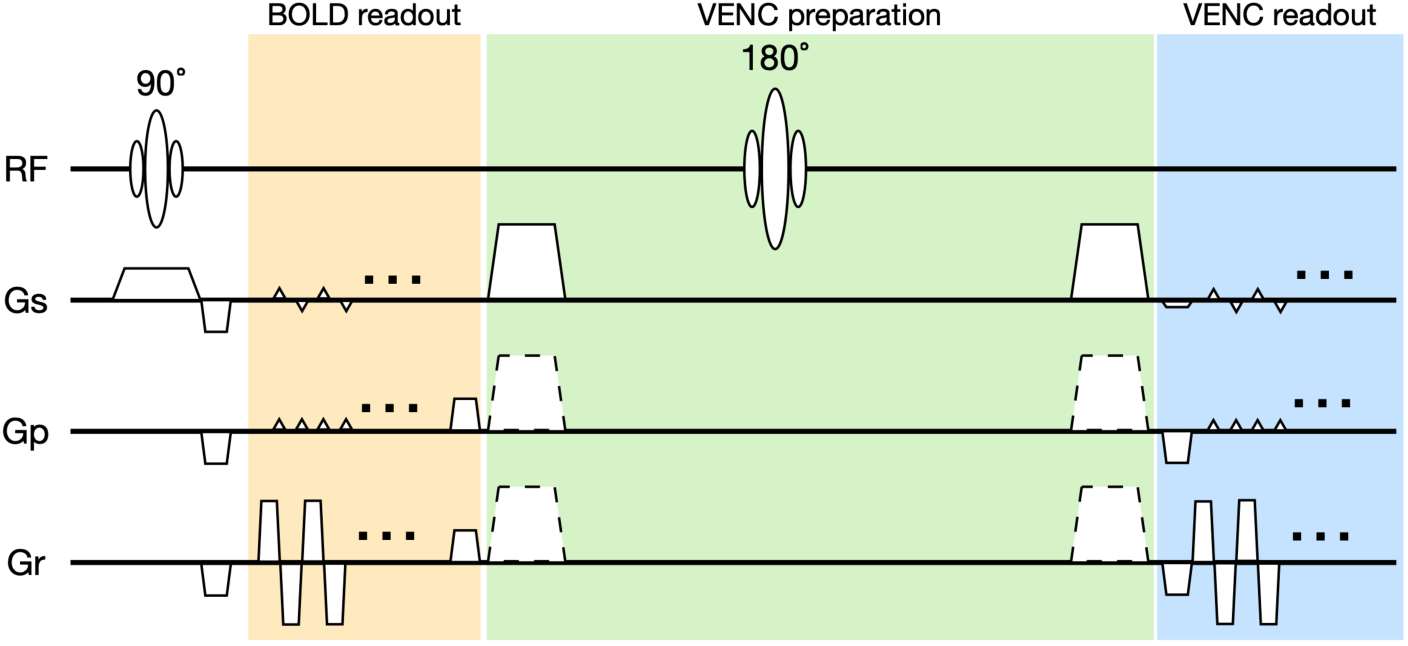
Pulse sequence diagram of the proposed BOLD-VENC sequence. The BOLD readout, VENC preparation module, and VENC readout are highlighted in yellow, green, and blue, respectively. Velocity-encoding gradients are applied along one of the slice (Gs), phase-encoding (Gp), or readout (Gr) directions. A relatively long echo time (TE) is used for the VENC readout to suppress signal contributions from blood vessels and enhance sensitivity to slow cerebrospinal fluid (CSF) flow.

### 2.2 Flow phantom experiments

To evaluate the accuracy and reliability of velocity measurement, we constructed a slow-flow phantom to evaluate slow-flow velocity measurements over a physiologically relevant velocity range. A plastic tube was used to simulate bulk CSF movement through CSF-rich anatomical spaces such as the fourth ventricle and subarachnoid space. To provide a stationary reference, the tube was wrapped around a spherical water phantom. As shown in **Error! Reference source not found.**, water entered the phantom from one end (#2), running through the tube and exited at the other end (#3). Flow was driven by an electronic syringe pump with a 50-mL syringe. The pump was programmed to generate mean flow velocities of 0.1, 0.2, 0.5, and 1.0 mm/s. To further evaluate the effect of inflow on the fMRI signal, we performed an additional experiment in which the syringe pump remained off during the first minute of image acquisition and was subsequently switched on to generate a flow velocity of 0.5 mm/s.

### 2.3 Participants and MR acquisitions

To validate BOLD-VENC method, data were acquired in vivo from five healthy adults with the experimental protocol approved by the Institutional Review Board. Prior to participation, each individual provided written informed consent. MR images were acquired using a Siemens 3T Prisma scanner (Siemens Healthcare, Erlangen, Germany) and a 32-channel head coil at the Biomedical Research Imaging Center (BRIC) at the University of North Carolina at Chapel Hill.

### 2.4 Paced breathing task

Because respiration is known to modulate CSF flow in the ventricular system (Dreha-Kulaczewski et al., 2015; Kollmeier et al., 2022), two participants performed a paced breathing task consisting of alternating 5-s inhalation and 5-s exhalation for 5 minutes in the scanner. However, respiration can also induce B0 fluctuations, leading to respiration-synchronized phase changes (Van De Moortele et al., 2002) that are unrelated to CSF flow. To control for the effects of B0 fluctuations on the measured phase signal, the VENC gradient was disabled during the first 30 s of the scan and enabled for the remaining 4.5 minutes. Velocity encoding was applied along one of the three orthogonal gradient directions in each session. A total of three sessions were acquired for each participant, with a different VENC encoding direction used in each session.

### 2.5 Breath-holding task

Inspiration has been shown to enhance CSF flow, whereas breath holding suppresses it (Dreha-Kulaczewski et al., 2015). To validate the VENC imaging technique, we employed a breathing paradigm consisting of 18 s of paced breathing followed by 15 s of breath holding, as illustrated in Figure 4a. During the paced breathing period, participants alternated 3-s inhalation and 3-s exhalation. Each breath-holding period was followed by 21 s of normal breathing before the next trial. The task consisted of five repetitions of this breathing paradigm and was preceded by the acquisition of five b0 images before task onset. The total acquisition time for each session was 5 minutes and 15 seconds. As in the paced breathing experiment (Section 2.4), velocity encoding was applied separately along the three orthogonal gradient directions in three imaging sessions. Throughout the experiment, respiratory and cardiac physiological signals were continuously recorded using the scanner’s built-in physiological monitoring system (Siemens Healthcare, Erlangen, Germany).

### 2.7 Visual stimulation task

To evaluate the relationship between localized BOLD activity and regional CSF dynamics, a high-contrast visual stimulation task was presented to participants, as illustrated in Figure 6a. The checkerboard subtended 20° of visual angle and was generated from 20 evenly distributed radial wedges (18° each) and eight concentric rings of equal width. The checkerboard contrast reversed at 12 Hz, corresponding to a contrast inversion every 1/24 s. Each stimulation block consisted of 15 s of checkerboard presentation, followed by 30–33 s of a uniform gray background with a central fixation cross. A 3-s temporal jitter was introduced to reduce anticipation effects. Six stimulation blocks were presented in each imaging session. The total acquisition time per session was 4 minutes and 51 seconds, including the acquisition of 5 b0 images before task onset. As in the breathing experiments, three imaging sessions were performed with VENC applied separately along each of the three orthogonal gradient directions. The event-related responses were analyzed using FSL FEAT (Woolrich et al., 2004). Statistical maps were obtained using an uncorrected threshold of p < 0.05 and cluster-wise correction with p < 0.05. The individual corrected activation maps were used as functional masks for the cross-correlation analysis described in Section 2.12.

### 2.8 Velocity analysis

Before velocity quantification, the background phase was removed from the phase images. In this study, background phase correction was performed in two steps: 1) k-space filtering using a two-dimensional (2D) Hamming filter and 2) 2D second-order polynomial fitting. This two-step approach was adopted instead of polynomial fitting alone to mitigate the effects of phase singularities. Phase singularities are characterized by abrupt spatial phase variations and may arise from magnetic field inhomogeneity or multi-channel coil combination. Such rapid phase variations cannot be adequately modeled by a low-order polynomial. By first applying a Hamming filter in k-space, the high-spatial-frequency phase components associated with these singularities were largely suppressed, allowing the remaining slowly varying background phase to be accurately estimated using polynomial fitting.

In the first step, the static background phase was estimated from the b0 images. The five b0 images acquired at the beginning of each scan were averaged and Fourier-transformed into k-space. After applying a 2D Hamming filter in k-space, the phase of the filtered image was regarded as the static background phase and subtracted from all image slices and time points. In the second step, the residual dynamic background phase was estimated separately for each simultaneous multi-slice (SMS) slice group and each time point using 2D second-order polynomial fitting. To prevent the velocity-encoded phase at CSF-rich regions from biasing the polynomial fit, voxels with high signal intensity in the VENC magnitude images were excluded from the fitting procedure. High-intensity voxels were defined as those with signal intensities exceeding 10% of the maximum image intensity.

### 2.9 Basic image processing

Reconstructed images were denoised using the NORDIC method, specifically by employing the "NIFTI_NORDIC" function in the NORDIC toolbox. After denoising, velocity analysis was performed prior to motion correction. The BOLD image series were then motion corrected using FSL (Jenkinson et al., 2012). The estimated rigid-body transformations and all subsequent image processing steps were also applied to the VENC and velocity images. For data acquired during the visual stimulation task, slice timing correction was performed using Hamming-windowed sinc interpolation. For data acquired during the breathing tasks, slice timing correction was not performed because the images were subsequently retrospectively gated to the respiratory and cardiac cycles (see Section 2.10). Finally, distortion correction was performed using the topup command in FSL.

### 2.10 Retrospective gating

Using the recorded physiological signals, the peaks of the respiratory waveform were identified to retrospectively reconstruct respiration-gated velocity responses. In this study, the respiration-gated responses were divided into 10 temporal frames (1 s per frame) centered on the respiratory peaks. Because the SMS groups were acquired every 250 ms, four consecutive SMS groups were assigned to each temporal frame. Likewise, the cardiac-gated responses were divided into four temporal frames (0.25 s per frame). To accurately determine the acquisition timing of every voxel, an acquisition timing image series was generated. The same rigid-body motion correction and distortion correction that were applied to the VENC images were also applied to the acquisition timing image series using nearest-neighbor interpolation to preserve the discrete temporal labels.

### 2.11 Velocity vector maps

To visualize the CSF flow direction, representative in-plane velocity vector maps were reconstructed from the two velocity components lying within the displayed imaging plane. After retrospective respiratory gating (Section 2.10), the velocity images corresponding to representative respiratory phases were selected. The vector field was down-sampled by a factor of three in both spatial dimensions for visualization, and the vector magnitude was calculated as the Euclidean norm of the two in-plane velocity components. Vectors with magnitudes below the 30th percentile were omitted to reduce visual clutter. The velocity vectors were displayed as arrows color-coded using an HSV color wheel, in which the hue represented the in-plane flow direction, whereas the value (brightness) was proportional to the velocity. Arrow orientation and length also indicated the flow direction and relative velocity, respectively.

### 2.12 Cross-correlation

To investigate the relationship between regional BOLD activity and CSF dynamics, cross-correlation analyses were performed between the gray-matter BOLD signal and either CSF inflow or CSF velocity. The gray-matter mask was generated from FreeSurfer segmentation in each participant’s T1-weighted (T1w) image space. For signal extraction from the fourth ventricle, only 4^th^-ventricle voxels within the bottom slice were analyzed, because the inflow effect attenuated substantially beyond the second slice. The fourth-ventricle mask was initially generated from the individual FreeSurfer segmentation and then eroded by one voxel using the FSL command fslmaths to reduce partial-volume effects. Both the gray-matter and fourth-ventricle masks were subsequently coregistered to the EPI space.

Region-of-interest (ROI)-averaged BOLD time series were used as the seed time courses for the cross-correlation analysis. To reduce motion-related confounding, the six estimated rigid-body motion parameters were regressed out from the BOLD time series. The resulting BOLD time series were then upsampled by a factor of 10 and low-pass filtered at 0.15 Hz. For CSF velocity analysis, velocity images acquired along the three orthogonal encoding directions were coregistered across sessions. The velocity magnitude was calculated as the Euclidean norm of the three velocity components. CSF velocity time series were processed in the same manner including up-sampling and low-pass filtering. The calculation of the cross-correlation between BOLD and CSF inflow is illustrated in Figure 5a. The negative derivative of the BOLD time series was cross-correlated with the CSF inflow time series over a range of temporal lags. For the analysis of BOLD and CSF velocity, the covariance between the BOLD signal and each of the three orthogonal velocity components was calculated separately and then summed. The resulting correlation coefficient was calculated as the total covariance divided by the product of the standard deviations of the BOLD and CSF velocity time series.

## 3. RESULTS

### 3.1 The phantom validation

To evaluate the accuracy and reliability of the proposed velocity measurement, the flow phantom was scanned with four different flow velocities (0.1, 0.2, 0.5, and 1.0 mm/s). The estimated velocity maps are shown in Figure 2b. The four flow tubes exhibited the expected flow directions and increasing velocity with increasing pumping rates, whereas the static water phantom remained close to zero velocity under all experimental conditions. The absence of measurable velocity in the static water phantom indicates that the background phase was effectively removed and that the proposed method introduced minimal measurement bias.

**Figure 2:**
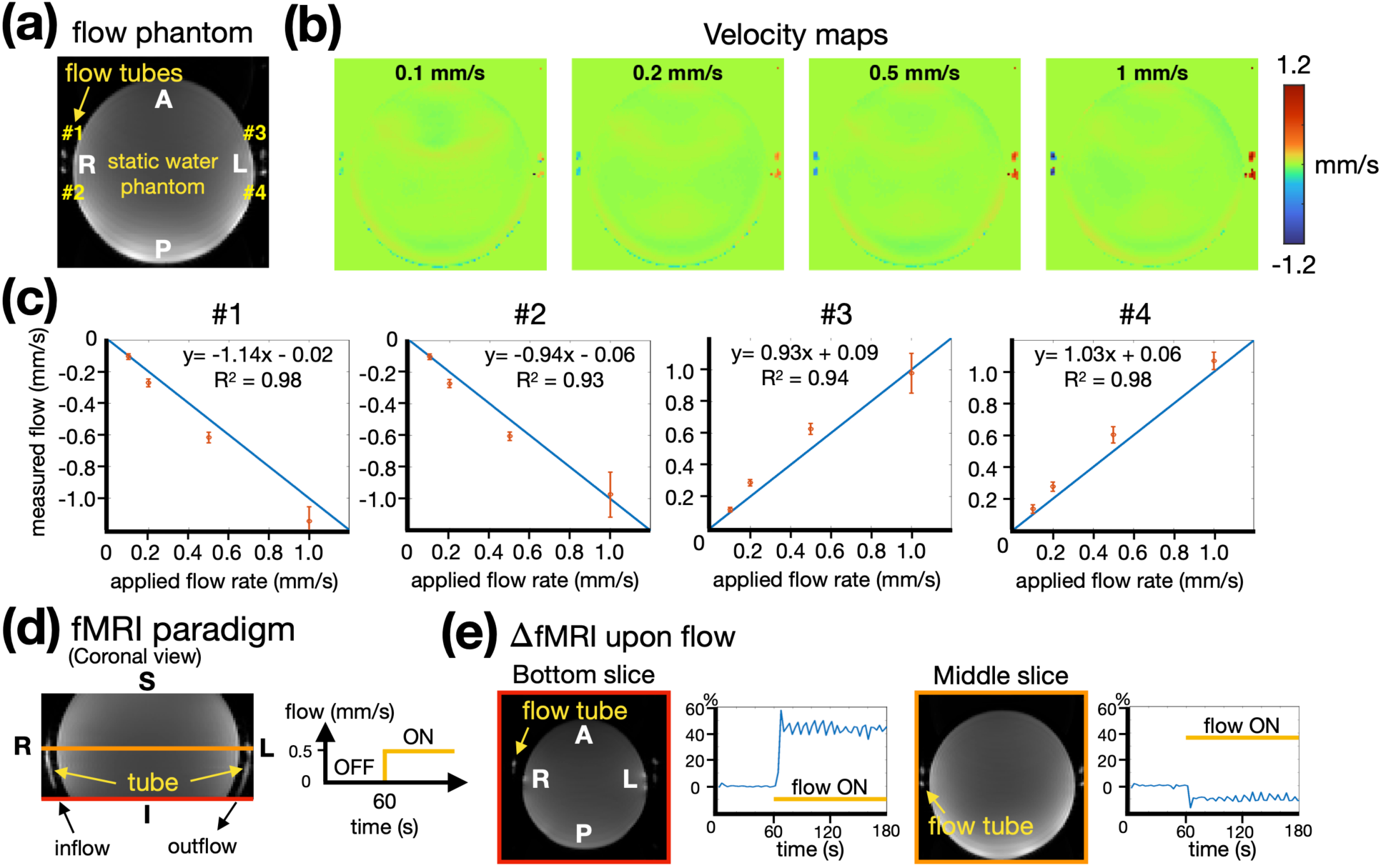
Validation of the BOLD-VENC sequence using a slow-flow phantom. (a) Axial MR image of the flow phantom. A single flow tube was wrapped around a static water phantom, producing four tube cross-sections (#1–#4). (b) Velocity maps acquired with the applied flow rate of 0.1, 0.2, 0.5, and 1.0 mm/s. The static water phantom remained close to zero velocity. (c) Linear relationship between the measured and applied flow velocities at the four tube cross-sections. Error bars represent the standard deviation across repeated measurements. (d) Experimental paradigm for evaluating the inflow sensitivity of the BOLD-VENC sequence. Water flow was switched from 0 to 0.5 mm/s at 60 s. (e) Time courses of the percentage fMRI signal change following flow onset. The signal increased in the bottom slice, whereas it decreased in the middle slice.

Quantitative analysis demonstrated general agreement between the measured and prescribed flow velocities as shown in Figure 2c. Across all four tube cross sections, the measured velocities exhibited strong linear relationships with the applied flow rates (R^2^ = 0.93–0.98), with regression slopes ranging from 0.93 to 1.14. Furthermore, the measurement variability remained small as indicated by the narrow error bars, demonstrating high measurement reliability. Notably, flow velocities as low as 0.1 mm/s were reliably detected, supporting the sensitivity of the BOLD-VENC sequence for very slow fluid motion.

To examine whether the proposed sequence remained sensitive to inflow despite the relatively long repetition time (TR = 3s) and a 90° excitation pulse, a flow-switching experiment was performed as illustrated in Figure 2d. At the bottom imaging slice (highlighted in red), where fresh spins entered the imaging volume, the fMRI signal increased following the onset of flow and remained significantly elevated throughout the flowing period (p < 0.001, Figure 2e). In contrast, the corresponding signal measured in a middle slice (highlighted in orange) decreased after flow onset. This reduction is consistent with inflow-related saturation of spins that had already experienced previous RF excitations, supporting that the signal enhancement observed in the bottom slice primarily originated from the inflow effect rather than from changes in bulk water properties. These results demonstrate that the proposed BOLD-VENC sequence preserves sensitivity to inflow at the bottom slice.

### 3.2 Paced Breathing

To characterize respiration-driven CSF dynamics, retrospective respiratory gating was applied to reconstruct velocity maps within a respiratory cycle. Representative respiration-gated velocity maps are shown in Figure 3a. The temporal characteristics of the velocity response were spatially heterogeneous. In particular, the fourth ventricle exhibited substantially higher flow velocities than the lateral ventricles and SAS. In addition, as shown in Figure 3b, the respiration-gated velocity profile in the fourth ventricle showed a pronounced increase during inhalation followed by a decrease during exhalation. The lateral ventricles exhibited a delayed and smaller-amplitude response. In contrast, the left and right peri-Sylvian SAS displayed nearly symmetric temporal profiles.

**Figure 3:**
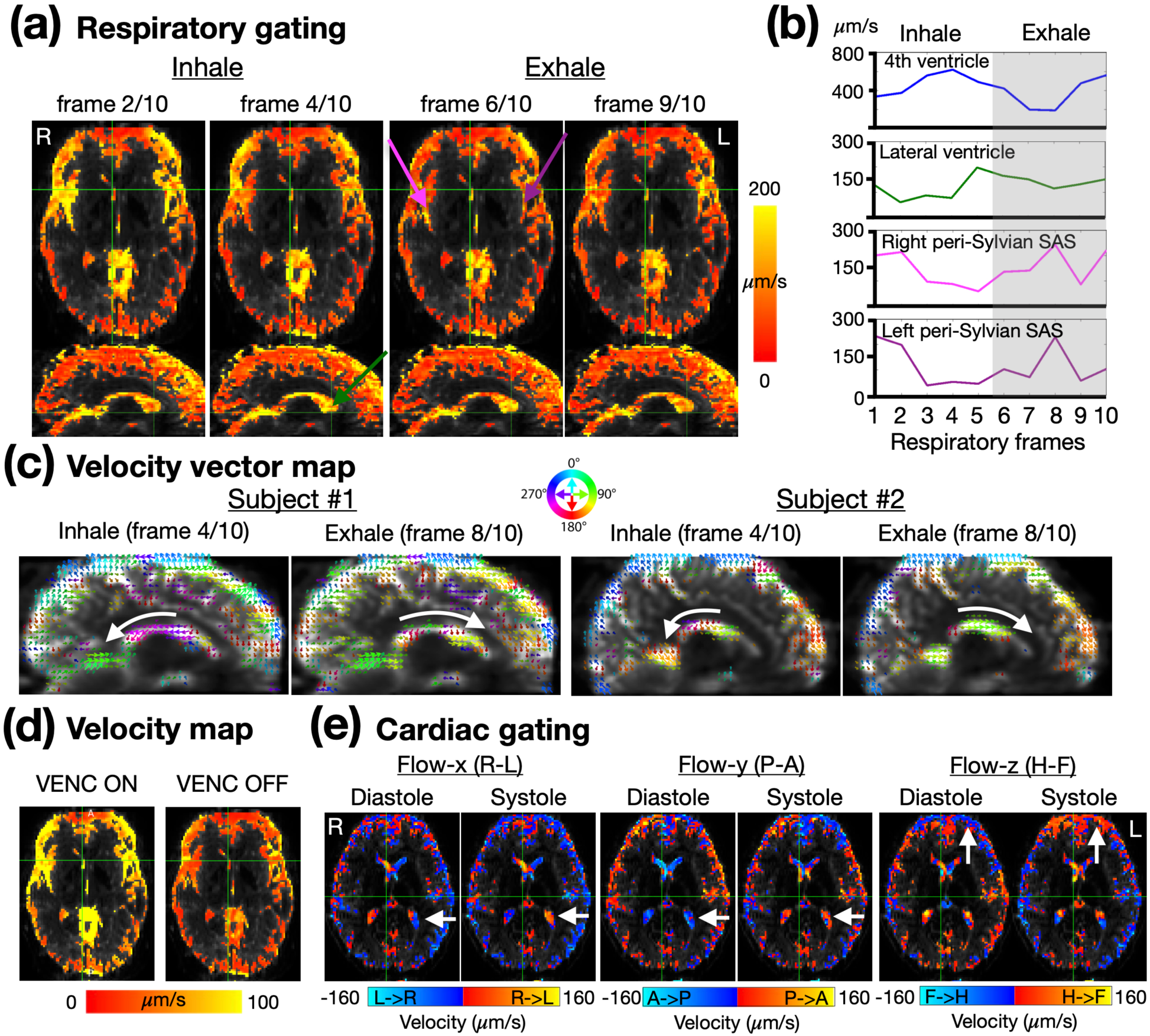
Respiratory- and cardiac-gated CSF velocity measurements. (a) Representative respiration-gated velocity maps. Frame 2 and 4 during inhalation and frames 6 and 9 during exhalation are shown. (b) CSF velocity profiles measured in the fourth ventricle, lateral ventricle, and bilateral peri-Sylvian SAS. (c) Velocity vector maps from two participants during inhalation and exhalation. The direction of CSF flow reversed between the two respiratory phases. (d) Velocity maps averaged across the respiratory cycle with VENC gradients enabled and disabled. (e) Cardiac-gated velocity maps reconstructed in three orthogonal directions during diastole and systole.

Velocity vector maps further demonstrated that respiration modulated not only the magnitude but also the direction of CSF flow (see Figure 3c). During inhalation, CSF within the lateral ventricles predominantly flowed from anterior to posterior, whereas the flow direction reversed during exhalation. To exclude the possibility that the measured velocities resulted from respiration-induced B0 fluctuations, the respiratory-gating analysis was repeated with the VENC gradients disabled. Figure 3d shows the velocity maps averaged across the respiratory cycle. The measured velocities were markedly reduced when VENC was turned off, confirming that the observed respiration-related velocity changes originated predominantly from velocity encoding rather than respiration-induced phase fluctuations.

Retrospective cardiac gating was also performed to evaluate cardiac-driven CSF motion (Figure 3e). Although the temporal resolution of 0.25 s was only marginally sufficient to resolve the rapid dynamics within a cardiac cycle, the reconstructed velocity maps revealed clear cardiac-driven flow patterns in all three orthogonal directions. The observed flow directions were in good qualitative agreement with the recently reported whole-brain CSF velocity maps obtained using slow-flow phase-contrast MRI (Dong et al., 2025), further supporting the validity of the proposed BOLD-VENC sequence for measuring CSF dynamics.

### 3.3 The breath holding

To further investigate CSF dynamics during respiratory challenges, participants performed a breath-holding task consisting of paced breathing, a 15-s breath-hold period, and recovery with normal breathing (see Figure 4a). The statistical maps of CSF flow velocity are shown in Figure 4b. Relative to normal breathing, inhalation induced a significant increase in CSF flow velocity, with the strongest responses observed in the lateral ventricles. Additional velocity increases were also observed in several cortical SAS. In contrast, breath holding produced a significant reduction in CSF flow velocity relative to normal breathing. Negative velocity responses were predominantly localized to the ventricular system, including the lateral ventricles and the third ventricle, indicating a marked attenuation of respiration-driven CSF motion during breath holding. These findings demonstrate that CSF flow velocity increases during inhalation but decreases during breath holding, consistent with previous reports of respiration-driven CSF flow dynamics (Dreha-Kulaczewski et al., 2015).

**Figure 4:**
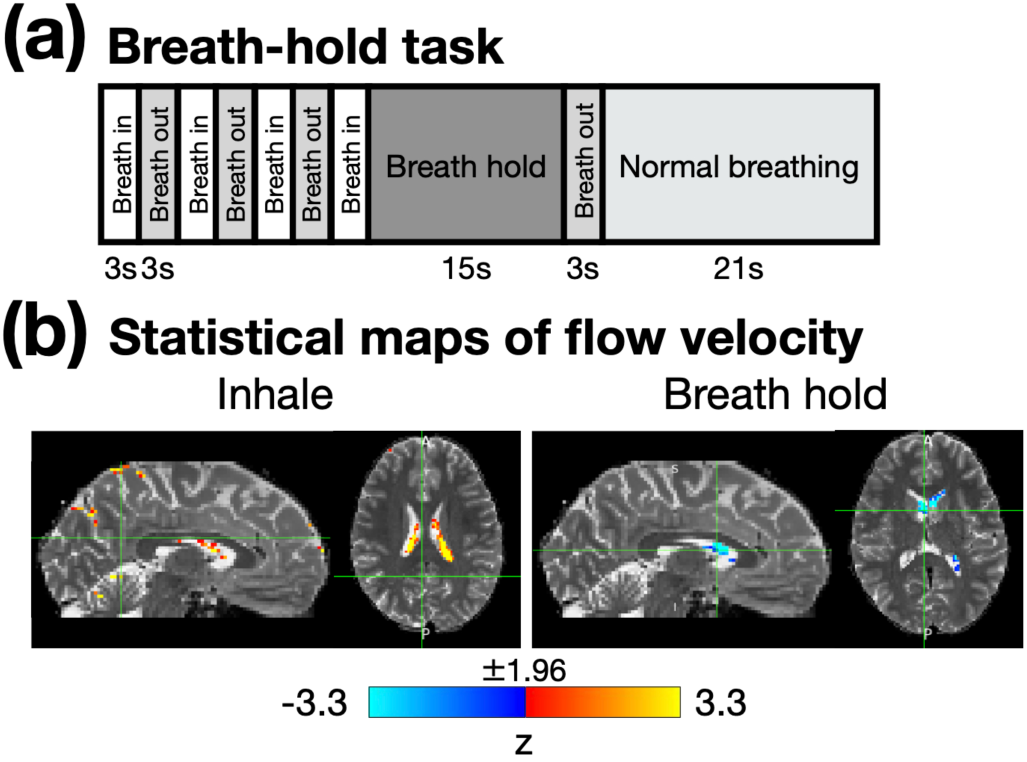
Breath-holding modulation of CSF flow velocity. (a) Breath-holding paradigm. (b) Representative statistical maps of CSF flow velocity from a participant. Z scores are color-coded as indicated by the color bar.

### 3.4 Coupling between CSF flow and global BOLD activity

To investigate the temporal relationship between gray-matter BOLD activity and CSF inflow at 4^th^ ventricle, the data acquired during the paced breathing and breath-holding tasks were analyzed using the pipeline illustrated in Figure 5a. The temporal derivative of the gray-matter BOLD signal was multiplied by −1 and shifted across a range of temporal lags prior to the cross-correlation analysis. Group-averaged cross-correlation functions (N = 5) revealed a robust positive association between the fourth-ventricle inflow signal and the negative derivative of the gray-matter BOLD signal, with the correlation peaking at a lag of 0.9 s (Figure 5b). The observed peak lag is consistent with previous reports showing that CSF inflow follows changes in the global BOLD signal with a short temporal delay (Fultz et al., 2019; Gu et al., 2022).

**Figure 5:**
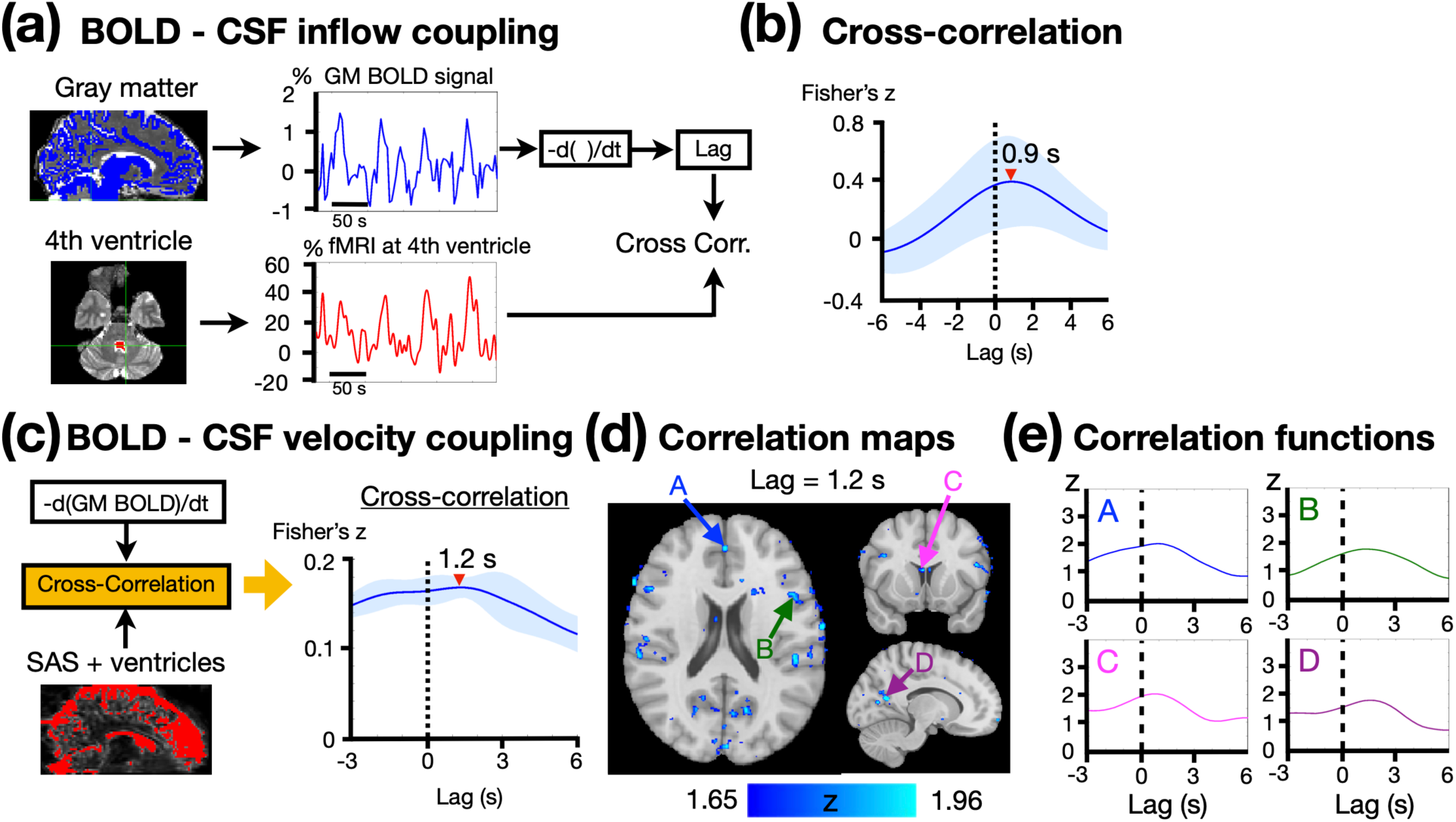
Coupling between gray-matter BOLD activity and CSF dynamics during the paced breathing and breath-holding tasks. (a) Schematic of the cross-correlation analysis between the negative derivative of the gray-matter BOLD signal and the CSF inflow at fourth ventricle. (b) Group-averaged (N = 5) cross-correlation function with the peak correlation at the lag of 0.9 s. Shaded regions indicate the standard error across participants. (c) Group-averaged cross-correlation function between ROI-averaged CSF flow velocity and gray-matter-averaged BOLD signal. The CSF ROIs include ventricles and cortical SAS. The peak correlation occurred at a lag of 1.2 s. (d) Group correlation map at the optimal lag of 1.2 s. Voxels were displayed only if they satisfied both the voxel-wise significance criterion for the peak correlation (p < 0.1) and the group-level significance criterion (t > 2.78, corresponding to p < 0.05). (e) Cross-correlation functions extracted from four regions of interest (A–D), demonstrating consistent temporal profiles with peak correlations occurring near a lag of 1.2 s.

We next examined the relationship between gray-matter BOLD activity and whole-brain CSF flow velocity measured with BOLD-VENC sequence. In Figure 5c, the time courses of flow velocity were averaged across SAS and ventricles prior to the correlation analysis. Similar to the inflow signal, the group-averaged CSF flow velocity exhibited a positive correlation with the negative derivative of the gray-matter BOLD signal, reaching a peak at a lag of 1.2 s (N = 5). Correlation maps were generated at this optimal lag and thresholded to retain only voxels that satisfied both a voxel-wise criterion (z of peak correlation > 1.65, corresponding to p < 0.1) and a group-level criterion (t > 2.78, corresponding to p < 0.05). The resulting maps revealed significant BOLD–velocity coupling in multiple ventricular and cortical SAS throughout the brain (Figure 5d). The correlation functions extracted from four regions of interest (A–D) demonstrated consistent temporal profiles, with peak correlations consistently occurring near a lag of 1.2 s, despite minor regional variations (Figure 5e). These results suggest that global BOLD fluctuations are temporally coupled with widespread CSF flow dynamics.

### 3.5 Coupling between CSF velocity and localized BOLD activity

To evaluate whether BOLD-VENC can detect task-evoked CSF dynamics, participants performed a visual stimulation task consisting of alternating 15-s checkerboard stimulation and 30–33 s fixation periods (see details in Section 2.7). Robust visual BOLD activation was consistently observed in all five participants, with activation localized primarily to the posterior calcarine sulcus and surrounding occipital visual cortex (Figure 6b). The group activation maps in the right-most panel demonstrated highly reproducible activation across subjects, confirming reliable detection of the visual response. For each participant, the corrected visual activation map (p < 0.05) was used to define a subject-specific ROI. The BOLD signal was averaged within the activation mask, and the negative derivative of the averaged BOLD signal was used as the seed time course for the BOLD-CSF velocity correlation analysis. Correlation maps were then computed using the optimal lag of 1.2 s, identified from the global BOLD-CSF velocity analysis during the respiratory tasks. The correlation maps were thresholded using the same criteria as those used in the resting-state cross-correlation analysis (Section 3.4). Significant BOLD-CSF velocity correlations were consistently observed across all participants (Figure 6c). The strongest correlations were localized to CSF spaces adjacent to the calcarine sulcus, with similar spatial distributions observed across subjects. Compared with the visual activation maps in Figure 6b, the significant correlations were generally located in the CSF spaces surrounding, rather than overlapping, the cortical activation sites. The group-averaged correlation map exhibited the same spatial pattern, demonstrating reproducible coupling between visually evoked BOLD activity and local CSF flow dynamics.

**Figure 6:**
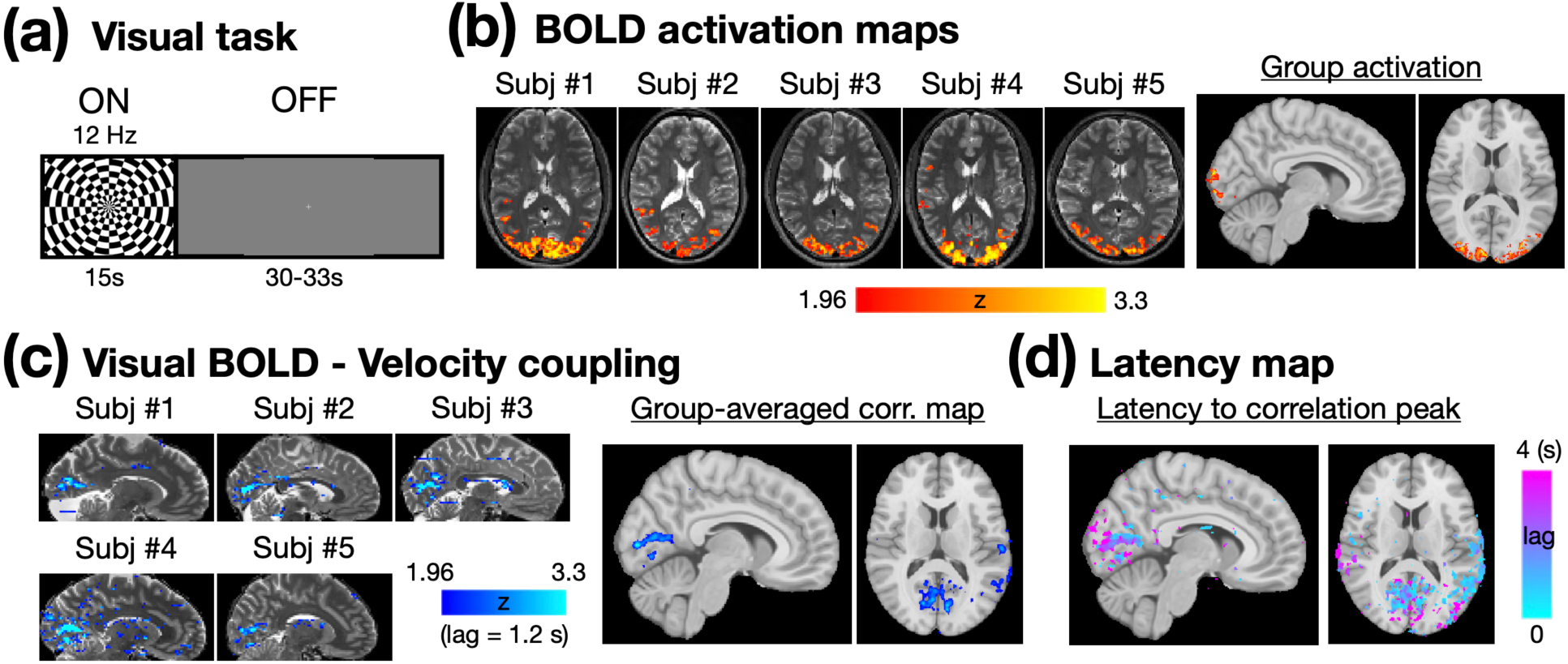
Task-evoked coupling between visual BOLD activity and CSF flow velocity. (a) Visual stimulation paradigm. (b) Individual and group BOLD activation maps during the visual task. (c) Individual and group correlation maps between BOLD signal and CSF flow velocity at the lag of 1.2 s. Correlation maps were thresholded using the same criteria as those used in breathing-related analyses (see Section 3.4). (d) Latency map showing the lag corresponding to the peak BOLD-velocity correlation at each voxel.

Using the cross-correlation functions at each voxel, a latency map can be generated by defining the latency of each voxel as the temporal lag yielding the maximum BOLD–velocity correlation (Figure 6d). Regions exhibiting short latencies were primarily distributed within the CSF spaces surrounding the visual cortex, closely corresponding to the regions showing significant BOLD-velocity correlations. These findings suggest that visually evoked neural activity induces rapid and spatially localized CSF flow responses in adjacent SASs.

## 4. DISCUSSION

In this study, we developed a single-shot BOLD–VENC sequence that combines a gradient-echo BOLD readout with a spin-echo velocity-encoded readout following the same RF excitation. The method provides brain-wide BOLD and slow-flow measurements with matched spatial coverage and closely aligned acquisition timing. The velocity measurement was validated in vitro, showing strong linear relationships between the measured and applied flow velocities across the tested range of 0.1-1.0 mm/s, with coefficients of determination of R^2^ = 0.93-0.98. The stationary water compartment remained close to zero, indicating that the applied background-phase correction did not introduce substantial velocity bias under the tested conditions. We further validated the VENC implementation in vivo within the established physiology of CSF motion under respiratory and cardiac influence. In paced breathing conditions, CSF velocity increased during inhalation with spatially varying responses across the fourth ventricle, lateral ventricles, and cortical subarachnoid spaces. Reversals in flow direction between inhalation and exhalation were also observed. These findings are consistent with previous reports that respiratory changes in intrathoracic pressure are transmitted through the venous system and contribute to craniospinal CSF displacement (Dreha-Kulaczewski et al., 2015; Kollmeier et al., 2022). When the VENC gradients were disabled, respiration-synchronized phase changes were markedly reduced, indicating that the predominant signal arose from velocity encoding rather than respiration-induced B0 fluctuations alone. Retrospective cardiac gating additionally revealed cardiac-phase-dependent flow patterns in all three orthogonal directions. Although the 0.25-s temporal resolution provided only a coarse representation of the cardiac cycle, the observed directions were in agreement with recent whole-brain slow-flow phase-contrast measurements (Dong et al., 2025). The reduction in ventricular velocity during breath holding was likewise consistent with the attenuation of respiration-driven CSF motion reported previously (Dreha-Kulaczewski et al., 2015). Taken together, in vitro and in vivo experiments support the validity of the velocity-encoded readout for measuring slow CSF motion.

We next used the combined acquisition to test whether regional CSF velocity changes were temporally related to simultaneously measured BOLD fluctuations. Previous inflow-sensitive fMRI studies identified coupling between the negative derivative of the global BOLD signal and CSF inflow at the fourth ventricle or craniospinal junction (Fultz et al., 2019; Han et al., 2021; Gu et al., 2022). In this study, this relationship was reproduced for fourth-ventricle inflow, with a peak correlation at approximately 0.9 s. Note that the global BOLD signal was also associated with velocity changes distributed across ventricular and cortical SAS, with a similar peak delay of approximately 1.2 s. The close agreement between these peak delays suggests that widespread CSF velocity fluctuations are driven by the same physiological process underlying the inflow at fourth ventricle. This interpretation is consistent with the model proposed by (Fultz et al., 2019), in which changes in cerebral blood volume are accompanied by compensatory CSF displacement under the constraint of a nearly fixed intracranial volume. Our results extend this framework by demonstrating that BOLD-associated CSF motion can be localized across multiple intracranial regions rather than being limited to a single inflow site. Because the measurements were obtained during paced-breathing and breath-holding conditions, in which respiratory pressure changes, arterial CO₂, autonomic activity, cerebral blood flow, and blood volume change concurrently, the observed coupling likely reflects contributions from both hemodynamic volume displacement and shared respiratory or autonomic modulation. A related strategy using pseudo-continuous arterial spin labeling (pCASL) also found coupling between cerebral hemodynamics and CSF motion, including a positive association between CBF and CSF pulsation during breath holding (Im et al., 2025). The proposed BOLD-VENC sequence additionally provides simultaneous measurements of CSF velocity magnitude and direction, enabling localization of BOLD-CSF coupling throughout the brain.

The visual-stimulation experiment further demonstrated a regional component of BOLD-CSF coupling. Velocity changes were localized primarily to subarachnoid spaces surrounding, rather than overlapping, the activated visual cortex. This spatial relationship suggests that local hemodynamic expansion of the activated cortex may mechanically displace CSF in nearby spaces, consistent with the volume-compensation mechanism proposed for global BOLD-CSF coupling and with previous visual-stimulation studies linking evoked hemodynamic responses to CSF inflow (Fultz et al., 2019; Williams et al., 2023) while not implying a direct causal relationship between neural activity and CSF motion (Logothetis et al., 2001). The corresponding latency map provides additional information about the temporal organization of this local response. Significant BOLD-CSF coupling was observed mainly in and around the activated visual cortex rather than in distant regions, with shorter latencies in areas surrounding the visual cortex. Given that stimulus-evoked neural activity provides a spatially localized driver of the vascular response, this pattern is consistent with a locally initiated vascular-CSF response that spreads to nearby regions. However, generalizing this interpretation of latency to paced-breathing or breath-holding experiments requires caution. Unlike visual stimulation, these respiratory tasks induce widespread physiological changes through multiple pathways. These processes can affect BOLD signals and CSF motion simultaneously but with different temporal responses across brain regions. Therefore, under respiratory challenges, regional BOLD-CSF latency may reflect the combined effects of multiple physiological processes rather than the propagation delay of a single vascular or CSF response.

Similar to the slow-flow-sensitized phase-contrast MRI method of (Dong et al., 2025), our BOLD-VENC sequence employs a long spin-echo TE (189 ms) to improve the specificity of CSF velocity measurements by suppressing signals from blood. At 3 T, the T2 of venous blood is ∼60 ms, leaving only ∼4% of the venous blood signal after T2 decay. In comparison, the slow-flow-sensitized phase-contrast MRI method at 7 T uses a TE of 80–88 ms. Using a representative venous blood T2 of 25 ms at 7T, the residual venous blood signal is ∼3% (Dong et al., 2025). Therefore, the residual venous blood signal is minimal and comparable between the two methods. For arterial blood, the T2 values at 3 T and 7 T are 150–170 ms and ∼68 ms respectively, resulting in a similar residual arterial blood signal of ∼30% after T2 decay in both implementations. In addition, arterial signal is expected to be further attenuated beyond that predicted by molecular diffusion alone. Arterial flow velocities are typically on the order of tens of centimeters per second, more than two orders of magnitude greater than the VENC (1.6 mm/s). Consequently, the bipolar velocity-encoding gradients are expected to generate substantial intravoxel phase dispersion and signal cancellation, thereby further reducing the contribution of arterial blood to the measured CSF velocity.

An important consideration for interpreting BOLD-VENC measurements, as well as other MRI-based approaches to glymphatic function, is that measurable CSF motion does not necessarily reflect fluid exchange with the interstitial space or the clearance of metabolic waste. BOLD-VENC measures coherent motion of MR-visible CSF but does not directly quantify CSF–interstitial fluid exchange or solute transport. Nevertheless, CSF motion represents an important component of brain fluid dynamics and may provide insight into the physiological forces that support fluid exchange and transport. Importantly, unlike conventional global BOLD-CSF coupling measures, BOLD-VENC enables simultaneous assessment of regional BOLD fluctuations and local CSF velocity, allowing BOLD-CSF coupling to be mapped across different ventricular and subarachnoid CSF spaces. This spatial information may be particularly valuable for clinical research because neurological disorders can affect vascular function and CSF pathways in a regionally heterogeneous manner. For example, reduced global BOLD-CSF coupling has been reported in Alzheimer’s disease (Han et al., 2021), but a global measure cannot determine whether this reduction arises from altered hemodynamic activity, impaired CSF motion, or changes associated with specific CSF pathways. By directly measuring the magnitude and direction of regional CSF velocity together with local BOLD activity, the BOLD-VENC sequence may help identify the spatial and physiological origins of altered BOLD-CSF coupling.

### Limitations

The current implementation has a spatial resolution of 2 mm isotropic. Although this resolution is sufficient to characterize CSF dynamics in the ventricles and larger SAS, higher spatial resolution will be needed to investigate smaller CSF spaces such as perivascular space (PVS). A spatial resolution of 1 × 1 × 2 mm^3^ is technically feasible with our sequence using an in-plane acceleration factor of 2, with minimal changes in TE and VENC. However, reducing the voxel volume from 8 to 2 mm^3^ substantially reduces the SNR, with an additional g-factor penalty associated with parallel imaging. Whether the phase precision and sensitivity are sufficient at this resolution requires further investigation.

Temporal resolution represents another limitation of the current implementation, which acquires one volume every 3 s for each velocity-encoding direction. Measurement of all three orthogonal velocity components currently requires repeated acquisitions, increasing the total scan time and limiting applications involving physiological events that cannot be reliably repeated. One potential strategy is to reduce the slice coverage and acquire the three velocity-encoding directions sequentially within a shorter time frame. Ultimately, the tradeoff between spatial resolution, temporal resolution, brain coverage, and SNR will depend on the specific scientific question.

Finally, as a proof-of-concept study, the present work was conducted in a relatively small cohort of healthy participants (N = 5), with the primary goal of demonstrating the feasibility of simultaneous BOLD and CSF velocity measurements. The reproducibility and interindividual variability of the observed regional BOLD-CSF coupling remain to be established. Future studies with larger cohorts and repeated measurements will be needed to evaluate the robustness of these findings and determine whether the observed relationships are preserved or altered across different physiological states, including rest and sleep, as well as across the lifespan and neurological disease.

## 5. CONCLUSION

This study demonstrates the feasibility of BOLD-VENC sequence for simultaneous measurement of brain-wide BOLD activity and spatially resolved slow CSF velocity at 3T. Beyond reproducing the established coupling between global BOLD fluctuations and fourth-ventricle CSF inflow, the BOLD-VENC sequence demonstrated BOLD-associated velocity changes across ventricles and cortical SAS. These findings extend BOLD-CSF coupling from a global measure toward a spatially resolved assessment of BOLD-CSF interactions. Such regional measurements may help determine whether impaired coupling in aging and neurological disorders is associated with diminished CSF motion in specific regions, altered local hemodynamic responses, or disruption of their temporal relationship.

## 6. Data and Code Availability

All data supporting the findings of this study will be made publicly available upon the publication of the manuscript on openneuro. This ensures that other researchers can replicate, verify, and build upon the work presented.

The computational code will also be made available to interested parties. Please note that the code is provided in its raw form and has not been optimized, cleaned, or extensively commented. While this may impact ease of use or adaptation, we believe it remains a valuable resource for those interested in understanding or extending our analytical methods.

## 7. Author contributions

M.W. contributed to manuscript preparation, including drafting portions of the manuscript. T.-H.C. contributed to the flow phantom experiments and data analysis. U.E. provided input on data analysis, interpretation, and presentation of the results. W.-T.C., the principal investigator, conceived and initiated the study and contributed to all aspects of the research, including methodology development and implementation, experimental design and execution, technical validation and troubleshooting, data analysis and interpretation, and manuscript preparation. W.-T.C. also had primary responsibility for project supervision, coordination, and integration of the collaborative efforts.

## REFERENCES

Da Mesquita, S., Louveau, A., Vaccari, A., Smirnov, I., Cornelison, R. C., Kingsmore, K. M., Contarino, C., Onengut-Gumuscu, S., Farber, E., Raper, D., Viar, K. E., Powell, R. D., Baker, W., Dabhi, N., Bai, R., Cao, R., Hu, S., Rich, S. S., Munson, J. M.,… Kipnis, J. (2018). Functional aspects of meningeal lymphatics in ageing and Alzheimer’s disease. Nature, 560(7717), 185–191. 10.1038/s41586-018-0368-8

Ding, X.-B., Wang, X.-X., Xia, D.-H., Liu, H., Tian, H.-Y., Fu, Y., Chen, Y.-K., Qin, C., Wang, J.-Q., Xiang, Z., Zhang, Z.-X., Cao, Q.-C., Wang, W., Li, J.-Y., Wu, E., Tang, B.-S., Ma, M.-M., Teng, J.-F., & Wang, X.-J. (2021). Impaired meningeal lymphatic drainage in patients with idiopathic Parkinson’s disease. Nature Medicine, 27(3), 411–418. 10.1038/s41591-020-01198-1

Dong, Z., Wang, F., Strom, A. K., Eckstein, K., Bachrata, B., Robinson, S. D., Rosen, B. R., Wald, L. L., Lewis, L. D., & Polimeni, J. R. (2025). Quantifying brain-wide cerebrospinal fluid flow dynamics using slow-flow-sensitized phase-contrast MRI. 10.1101/2025.03.22.644745

Dreha-Kulaczewski, S., Joseph, A. A., Merboldt, K.-D., Ludwig, H.-C., Gärtner, J., & Frahm, J. (2015). Inspiration Is the Major Regulator of Human CSF Flow. The Journal of Neuroscience, 35(6), 2485–2491. 10.1523/JNEUROSCI.3246-14.2015

Fultz, N. E., Bonmassar, G., Setsompop, K., Stickgold, R. A., Rosen, B. R., Polimeni, J. R., & Lewis, L. D. (2019). Coupled electrophysiological, hemodynamic, and cerebrospinal fluid oscillations in human sleep. Science, 366(6465), Article 6465. 10.1126/science.aax5440

Gu, Y., Han, F., Sainburg, L. E., Schade, M. M., Buxton, O. M., Duyn, J. H., & Liu, X. (2022). An orderly sequence of autonomic and neural events at transient arousal changes. NeuroImage, 264, 119720. 10.1016/j.neuroimage.2022.119720

Han, F., Chen, J., Belkin-Rosen, A., Gu, Y., Luo, L., Buxton, O. M., Liu, X., & the Alzheimer’s Disease Neuroimaging Initiative. (2021). Reduced coupling between cerebrospinal fluid flow and global brain activity is linked to Alzheimer disease–related pathology. PLOS Biology, 19(6), e3001233. 10.1371/journal.pbio.3001233

Iliff, J. J., Chen, M. J., Plog, B. A., Zeppenfeld, D. M., Soltero, M., Yang, L., Singh, I., Deane, R., & Nedergaard, M. (2014). Impairment of Glymphatic Pathway Function Promotes Tau Pathology after Traumatic Brain Injury. The Journal of Neuroscience, 34(49), 16180–16193. 10.1523/JNEUROSCI.3020-14.2014

Iliff, J. J., Wang, M., Liao, Y., Plogg, B. A., Peng, W., Gundersen, G. A., Benveniste, H., Vates, G. E., Deane, R., Goldman, S. A., Nagelhus, E. A., & Nedergaard, M. (2012). A Paravascular Pathway Facilitates CSF Flow Through the Brain Parenchyma and the Clearance of Interstitial Solutes, Including Amyloid β. Science Translational Medicine, 4(147). 10.1126/scitranslmed.3003748

Im, J.-G., Kim, J.-H., & Park, S.-H. (2025). Simultaneous measurement of cerebral blood flow and cerebrospinal fluid flow using pseudo-continuous arterial spin labeling. NeuroImage, 311, 121192. 10.1016/j.neuroimage.2025.121192

Jenkinson, M., Beckmann, C. F., Behrens, T. E., Woolrich, M. W., & Smith, S. M. (2012). Fsl. Neuroimage, 62(2), Article 2. 10.1016/j.neuroimage.2011.09.015 S1053-8119(11)01060-3 [pii]

Johanson, C. E., Duncan, J. A., Klinge, P. M., Brinker, T., Stopa, E. G., & Silverberg, G. D. (2008). Multiplicity of cerebrospinal fluid functions: New challenges in health and disease. Cerebrospinal Fluid Research, 5(1), 10. 10.1186/1743-8454-5-10

Kollmeier, J. M., Gürbüz-Reiss, L., Sahoo, P., Badura, S., Ellebracht, B., Keck, M., Gärtner, J., Ludwig, H.-C., Frahm, J., & Dreha-Kulaczewski, S. (2022). Deep breathing couples CSF and venous flow dynamics. Scientific Reports, 12(1), 2568. 10.1038/s41598-022-06361-x

Korbecki, A., Zimny, A., Podgórski, P., Sąsiadek, M., & Bladowska, J. (2019). Imaging of cerebrospinal fluid flow: Fundamentals, techniques, and clinical applications of phase-contrast magnetic resonance imaging. Polish Journal of Radiology, 84, 240–250. 10.5114/pjr.2019.86881

Kress, B. T., Iliff, J. J., Xia, M., Wang, M., Wei, H. S., Zeppenfeld, D., Xie, L., Kang, H., Xu, Q., Liew, J. A., Plog, B. A., Ding, F., Deane, R., & Nedergaard, M. (2014). Impairment of paravascular clearance pathways in the aging brain. Annals of Neurology, 76(6), 845– 861. 10.1002/ana.24271

Logothetis, N. K., Pauls, J., Augath, M., Trinath, T., & Oeltermann, A. (2001). Neurophysiological investigation of the basis of the fMRI signal. Nature, 412(6843), Article 6843. 10.1038/35084005 35084005 [pii]

Lotz, J., Meier, C., Leppert, A., & Galanski, M. (2002). Cardiovascular Flow Measurement with Phase-Contrast MR Imaging: Basic Facts and Implementation. RadioGraphics, 22(3), 651–671. 10.1148/radiographics.22.3.g02ma11651

Louveau, A., Smirnov, I., Keyes, T. J., Eccles, J. D., Rouhani, S. J., Peske, J. D., Derecki, N. C., Castle, D., Mandell, J. W., Lee, K. S., Harris, T. H., & Kipnis, J. (2015). Structural and functional features of central nervous system lymphatic vessels. Nature, 523(7560), 337–341. 10.1038/nature14432

Pelc, N. J., Bernstein, M. A., Shimakawa, A., & Glover, G. H. (1991). Encoding strategies for three-direction phase-contrast MR imaging of flow. Journal of Magnetic Resonance Imaging, 1(4), 405–413. 10.1002/jmri.1880010404

Ringstad, G., Valnes, L. M., Dale, A. M., Pripp, A. H., Vatnehol, S.-A. S., Emblem, K. E., Mardal, K.-A., & Eide, P. K. (2018). Brain-wide glymphatic enhancement and clearance in humans assessed with MRI. JCI Insight, 3(13), e121537. 10.1172/jci.insight.121537

Shen, M. D., Kim, S. H., McKinstry, R. C., Gu, H., Hazlett, H. C., Nordahl, C. W., Emerson, R. W., Shaw, D., Elison, J. T., Swanson, M. R., Fonov, V. S., Gerig, G., Dager, S. R., Botteron, K. N., Paterson, S., Schultz, R. T., Evans, A. C., Estes, A. M., Zwaigenbaum, L.,… Gu, H. (2017). Increased Extra-axial Cerebrospinal Fluid in High-Risk Infants Who Later Develop Autism. Biological Psychiatry, 82(3), 186–193. 10.1016/j.biopsych.2017.02.1095

Spijkerman, J. M., Petersen, E. T., Hendrikse, J., Luijten, P., & Zwanenburg, J. J. M. (2018). T 2 mapping of cerebrospinal fluid: 3 T versus 7 T. Magnetic Resonance Materials in Physics, Biology and Medicine, 31(3), 415–424. 10.1007/s10334-017-0659-3

Van De Moortele, P., Pfeuffer, J., Glover, G. H., Ugurbil, K., & Hu, X. (2002). Respiration-induced *B*_0_ fluctuations and their spatial distribution in the human brain at 7 Tesla. Magnetic Resonance in Medicine, 47(5), 888–895. 10.1002/mrm.10145

Williams, S. D., Setzer, B., Fultz, N. E., Valdiviezo, Z., Tacugue, N., Diamandis, Z., & Lewis, L. D. (2023). Neural activity induced by sensory stimulation can drive large-scale cerebrospinal fluid flow during wakefulness in humans. PLOS Biology, 21(3), e3002035. 10.1371/journal.pbio.3002035

Woolrich, M. W., Behrens, T. E. J., & Smith, S. M. (2004). Constrained linear basis sets for HRF modelling using Variational Bayes. NeuroImage, 21(4), Article 4. 10.1016/j.neuroimage.2003.12.024

Yamada, S., Miyazaki, M., Kanazawa, H., Higashi, M., Morohoshi, Y., Bluml, S., & McComb, J. G. (2008). Visualization of Cerebrospinal Fluid Movement with Spin Labeling at MR Imaging: Preliminary Results in Normal and Pathophysiologic Conditions. Radiology, 249(2), 644–652. 10.1148/radiol.2492071985

Zhang, W., Zhou, Y., Wang, J., Gong, X., Chen, Z., Zhang, X., Cai, J., Chen, S., Fang, L., Sun, J., & Lou, M. (2021). Glymphatic clearance function in patients with cerebral small vessel disease. NeuroImage, 238, 118257. 10.1016/j.neuroimage.2021.118257

Zhao, J. M., Clingman, C. S., Närväinen, M. J., Kauppinen, R. A., & Van Zijl, P. C. M. (2007). Oxygenation and hematocrit dependence of transverse relaxation rates of blood at 3T. Magnetic Resonance in Medicine, 58(3), 592–597. 10.1002/mrm.21342

